# Body weight at young adulthood and association with epigenetic aging and lifespan in the BXD murine family

**DOI:** 10.1101/791582

**Authors:** Jose Vladimir Sandoval-Sierra, Alexandra H. B. Helbing, Evan G. Williams, David G. Ashbrook, Suheeta Roy, Robert W. Williams, Khyobeni Mozhui

**Author notes:** Correspondence: Khyobeni Mozhui.

## Abstract

DNA methylation (DNAm) is shaped by genetic and environmental factors and modulated by aging. Here, we examine interrelations between epigenetic aging, body weight (BW), and lifespan in 12 inbred mouse strains from the BXD panel that exhibit over two-fold variation in longevity. Genome-wide DNAm was assayed in 70 liver specimens from mice ranging in age from 6 to 25 months that were maintained on normal chow or high fat diet (HFD). We defined subsets of CpG regions associated with age, BW at young adulthood, and strain-by-diet dependent life expectancy. The age associated differentially methylated CpG regions (age-DMRs) featured distinct genomic characteristics, with DNAm gains over time occurring in sites with high CpG density and low average methylation. CpG regions associated with BW were enriched in introns and generally showed lower methylation in mice with higher BW, and inversely correlated with gene expression such that mRNA was higher in mice with higher BW. Lifespan-associated regions featured no distinct genomic characteristics but were linked to genes involved in lifespan regulation, including the telomerase reverse transcriptase gene, *Tert*, which showed lower methylation and higher gene expression in long-lived strains. An epigenetic clock defined from the age-DMRs conveyed accelerated aging in mice belonging to strains with shorter lifespans. Both higher BW at young adulthood and HFD were associated with accelerated epigenetic aging. Our results highlight the age-accelerating effect of heavier body weight. Furthermore, the study demonstrates that the measure of epigenetic aging derived from age-DMRs can predict strain and diet-induced differences in lifespan.

## Introduction

In the past few years, the “epigenetic clock” based on DNA methylation (DNAm) has emerged as a robust and widely used biomarker of aging that perhaps surpasses telomere length assays in its accuracy and utility^1^. Often referred to as DNA methylation age (DNAmAge), the CpG based estimator of biological age comes in a few different versions for both humans and mice^2–8^. All these clocks share a common feature—they rely on the methylation status of preselected subsets of CpGs that are each assigned weights and are used collectively to estimate age. A critical question has been: are these DNAmAge clocks detecting changes that are purely a function of time, and therefore, correlates of chronological age? Or are they providing a measure of the intrinsic pace of biological aging that can be related to health, fitness, and life expectancy? Evidence from retrospective human epidemiological studies indicate that certain versions of the clock perform better at measuring biological aging and at predicting life expectancy. In general, a younger DNAmAge relative to chronological age conveys decelerated biological aging, and is associated with a lower risk of disease and increased longevity^4,9–12^.

In the mouse model, lifespan extending interventions such as calorie restriction (CR) and treatment with rapamycin, or strong genetic mutations that drastically reduce body weight (e.g., *Ames* and *Snell* dwarf mice), have been shown to significantly decelerate the epigenetic clock^5,7,8,13,14^. However, to date, most DNAmAge estimations in the mouse have been modeled on the canonical C57BL/6J (B6) background strain^5–8,13^. Similar to interindividual variation in humans, aging trajectories vary considerably between different mouse strains, and common DNA variants contribute to the pace of normal aging and longevity^15^. Whether the differential rates of epigenetic aging can discern normative lifespan differences between mouse strains has not yet been explored. Furthermore, if body weight reduction due to CR or dwarfing mutations can slow down the clock, then an open question is whether subtle differences in normal body weight can also have an impact on DNAmAge.

The BXD recombinant inbred family is the most deeply phenotyped mouse genetic reference panel, and has a long history in aging and longevity research^16–18^. Notably, the different BXD strains exhibit wide variation in natural life expectancy under normal laboratory conditions. Some strains have a mean life expectancy of less than 15 months (e.g., BXD13, BXD5), while other strains are expected to survive for well over 2 years (e.g., BXD19, BXD65)^16,18–20^. Longevity data in the BXDs has been collected since the 1980s, and lifespan data continues to be collected from the current, expanded BXD panel that consists of 150 inbred strains^18,20,21^. The BXDs are derived by crossing the two parental strains, B6 and DBA/2J (D2), and then inbreeding the F2s, or by advanced intercrosses to increase recombination^21^. Genomes of the BXDs therefore represent random recombinants of the B6 and D2 genomes, where each strain is a unique mosaic of homozygous B6 or D2 alleles. The D2 strain has been considered to have a more accelerated aging profile, and has a significantly shorter lifespan than B6^15,22^. Other age-associated parameters include rapid thymic involution^23^, quicker replicative senescence of hematopoietic stem cells^17^, and increased tail tendon breakage in D2 compared to B6^24^. Due to random assortment of independent gene variants, the progeny BXDs have a much greater range of variation in lifespan and aging traits^17,20^, and provide a unique population in which to dissect the interrelations between epigenetic aging and longevity.

Here, we leveraged the extensive longevity data generated for the BXD family^20^ to evaluate associations between body weight, epigenetic aging, and lifespan. We used affinity-capture enrichment with the methyl-CpG-binding domain protein (MBD), followed by deep sequencing (MBD-seq) to profile the aging liver methylome in 12 members of the BXD family^25^. To examine the impact of a common metabolic stressor, we also quantified the methylome on a subset of BXD mice maintained on a high fat diet (HFD), which decreases longevity by as much as 12%^20^. Our goal was to chart differentially methylated regions related to aging, and to strain differences in body weight and lifespan, and to examine correlation with gene expression. We computed a simple DNAm clock using the age-dependent CpG regions, and this conveyed strain differences in rates of epigenetic aging. Overall, our results show that both higher body weight and HFD can have an accelerating effect on epigenetic aging, and this can then be related to lifespan.

## Results

### BXD strain selection and lifespan and body weight characteristics

The present work is based on data collected from two separate cohorts of BXD mice. Lifespan data were collected from a group of female BXDs, referred to as the *“longevity cohort”*, that were allowed to age until mortality^20^. A parallel, identically treated and strain-matched group, the *“biospecimen cohort”,* was maintained for tissue collection at different ages, and liver samples were obtained from this. The BXD strains have wide range in natural lifespan when kept on *ad libitum* standard chow (control diet or CD)^20^. HFD reduced longevity, but with marked differences among strains^20^. For DNAm analysis, we selected 12 members of the BXD panel (including F1 hybrids) that were representative of the wide variation in lifespan, and for five of these, we included mice maintained on HFD (**Table 1; Fig 1a**). Ages at natural death from the *longevity cohort* are plotted in **Fig 1a**. BXD65 showed the most significant decline in lifespan on HFD.

**Fig. 1.**
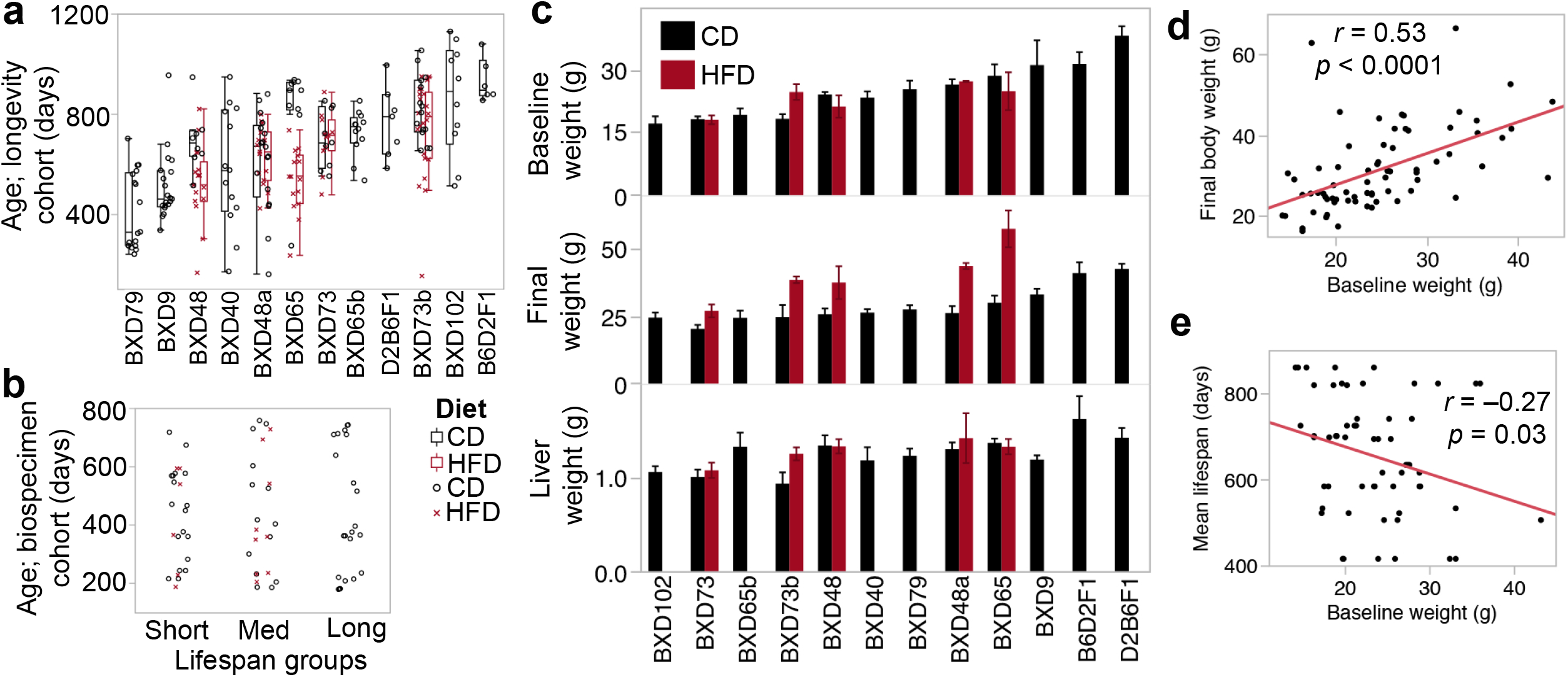
Age distribution and body weight characteristics. **(a)** Each point depicts a mouse that was allowed to age till mortality. There were a total of 225 mice for the 17 strain-by-diet groups on control diet (CD; black circles) or high fat diet (HFD; red crosses). **(b)** Each point depicts a mouse used for tissue collection and methylome assay. Age distribution (y-axis)is uniform across the three lifespan groups. **(c)** The bar plots show average body weight at young adulthood (baseline weight), and body and liver weight at final weighing (error bars are standard error). The baseline body weight (x-axis) was **(d)** a significant predictor of individual body weight at older age (y-axis), and **(e)** was negatively correlated with mean lifespan (y-axis) for the strain-by-diet group.

Based on lifespan averages, each strain-by-diet group was classified as short-lived (mean lifespan < 600 days), medium-lived (600-750 days), and long-lived (>800 days; no strain on HFD were in this group). Note that a strain classified as long-lived on CD may be classified as short-lived on HFD (**Table 1**). We then selected 70 liver specimens for the corresponding 17 strain-by-diet groups for DNAm assay. These specimens were chosen so that distribution of age at time of tissue collection was closely matched across the lifespan groups (**Fig. 1b;** individual level sample information in **Table S1**). We note that aside from three male cases for BXD102, B6D2F1, and D2B6F1 (**Table 1**), all liver specimens were from females. While the samples were not chosen on the basis of body or organ weight, there was significant strain variation in body weight when the mice were initially weighed at young adulthood (mean age of 134 ± 81 days), before introduction to HFD (**Fig. 1c**). We refer to this as baseline body weight or BW0. The long-lived F1s had higher body weights, and this hybrid vigor was apparent with or without the male cases. The final weight of mice (i.e., weight on day of sample collection) continued to show significant strain variation (**Fig. 1c**). Liver weight appeared fairly consistent across strains (**Fig. 1c**). There was no group difference in BW0 between the two diets (CD = 25 ± 7 g vs. HFD = 23 ± 5 g; *n* = 70). By final weighing, the group on HFD had become significantly heavier when compared to the full set of mice on CD (HFD = 41 ± 12 g vs. CD = 29 ± 8 g, p < 0.0001; n = 70), or when compared only to the matched strains on CD (HFD = 41 ± 12 g vs. CD = 26 ± 6 g, p < 0.0001; n = 34). The weight of the liver on HFD was slightly heavier but the effect was not statistically significant, likely due to the modest sample size for HFD (HFD = 1.29 ± 0.23 g vs. CD = 1.22 ± 0.23 g; p = 0.37, n = 34). At young adulthood, age was a significant predictor of BW0 (r = 0.55, p < 0.0001, n = 70). By the age at which samples were collected, the age of mice was no longer correlated with final body weight (r = 0.01) or with weight of liver (r = 0.12). Instead, BW0 remained a significant predictor of final body (**Fig. 1d**), and liver weight (r = 0.53, p < 0.0001, n = 70). When restricted to only the few HFD mice, BW0 was still a significant correlate of final body weight (r = 0.53, p = 0.05, *n* = 15), but not liver weight (r = 0.28, p = 0.30).

We next examined correlations between the individual weight measures from the *biospecimen cohort,* and strain-level lifespan from the *longevity cohort*. The F1s exhibited vigor in both longevity and body weight and including the F1s resulted in no significant correlation between weight and lifespan. After excluding the F1s, both BW0 and final weight showed inverse correlations with all the strain-level indicators of lifespan (mean, median, maximum, and minimum lifespan) (**Table S2**). This indicates that mice with higher body weight, even at younger age, are more likely to belong to strains with shorter lifespan (**Fig 1e**). This observation is consistent with the strong inverse correlation between body weight and longevity that is seen in the larger BXD panel^20^.

**Table 1.**
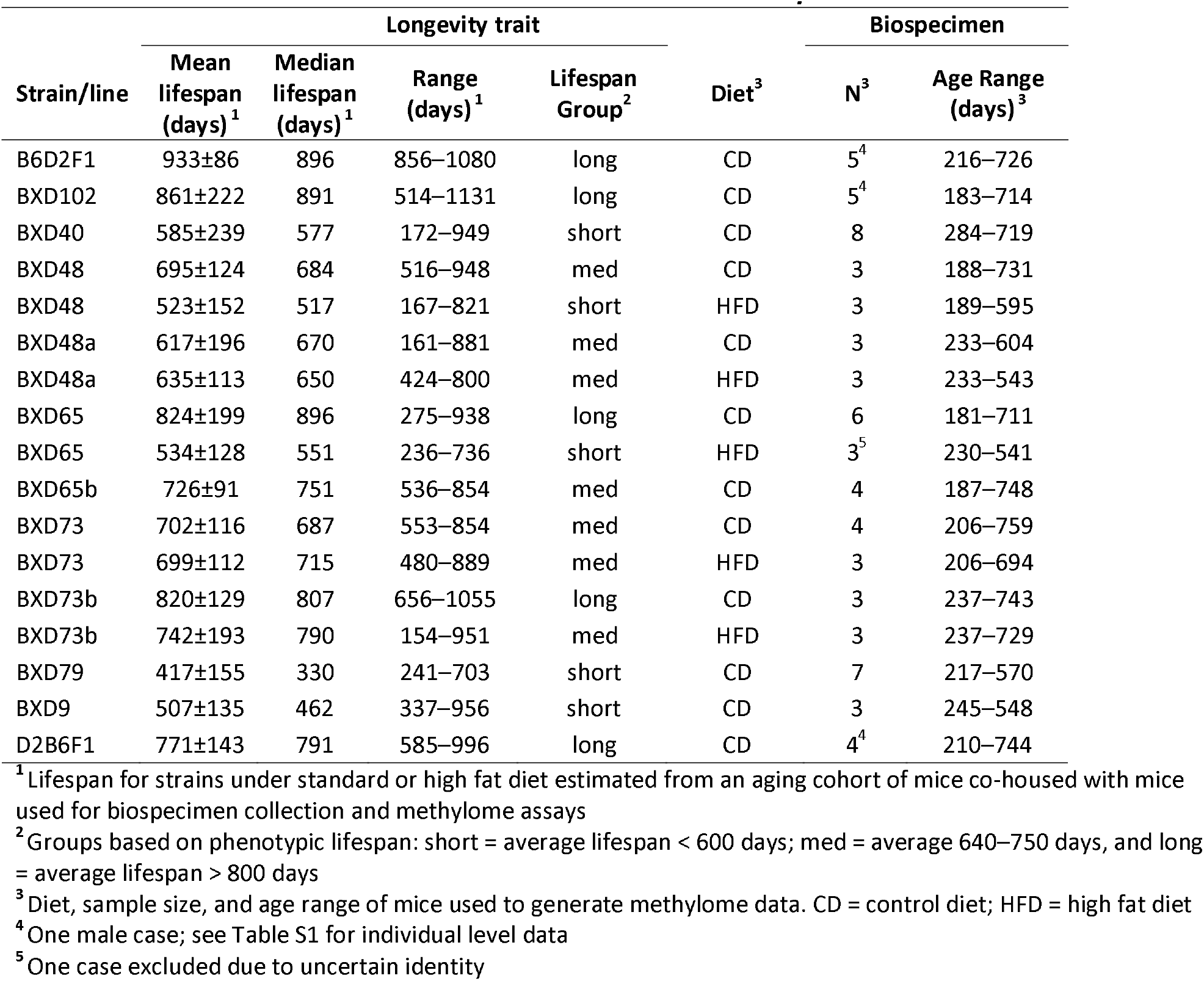
Characteristics of selected strains from the BXD family

| Strain/line | Longevity trait |  |  |  | Diet<sup>3</sup> | Biospecimen |  |
| --- | --- | --- | --- | --- | --- | --- | --- |
|  | Mean lifespan (days)<sup>1</sup> | Median lifespan (days)<sup>1</sup> | Range (days)<sup>1</sup> | Lifespan Group<sup>2</sup> |  | N<sup>3</sup> | Age Range (days)<sup>3</sup> |
| B6D2F1 | 933±86 | 896 | 856–1080 | long | CD | 5<sup>4</sup> | 216–726 |
| BXD102 | 861±222 | 891 | 514–1131 | long | CD | 5<sup>4</sup> | 183–714 |
| BXD40 | 585±239 | 577 | 172–949 | short | CD | 8 | 284–719 |
| BXD48 | 695±124 | 684 | 516–948 | med | CD | 3 | 188–731 |
| BXD48 | 523±152 | 517 | 167–821 | short | HFD | 3 | 189–595 |
| BXD48a | 617±196 | 670 | 161–881 | med | CD | 3 | 233–604 |
| BXD48a | 635±113 | 650 | 424–800 | med | HFD | 3 | 233–543 |
| BXD65 | 824±199 | 896 | 275–938 | long | CD | 6 | 181–711 |
| BXD65 | 534±128 | 551 | 236–736 | short | HFD | 3<sup>5</sup> | 230–541 |
| BXD65b | 726±91 | 751 | 536–854 | med | CD | 4 | 187–748 |
| BXD73 | 702±116 | 687 | 553–854 | med | CD | 4 | 206–759 |
| BXD73 | 699±112 | 715 | 480–889 | med | HFD | 3 | 206–694 |
| BXD73b | 820±129 | 807 | 656–1055 | long | CD | 3 | 237–743 |
| BXD73b | 742±193 | 790 | 154–951 | med | HFD | 3 | 237–729 |
| BXD79 | 417±155 | 330 | 241–703 | short | CD | 7 | 217–570 |
| BXD9 | 507±135 | 462 | 337–956 | short | CD | 3 | 245–548 |
| D2B6F1 | 771±143 | 791 | 585–996 | long | CD | 4<sup>4</sup> | 210–744 |

<sup>1</sup> Lifespan for strains under standard or high fat diet estimated from an aging cohort of mice co-housed with mice used for biospecimen collection and methylome assays
<sup>2</sup> Groups based on phenotypic lifespan: short = average lifespan < 600 days; med = average 640–750 days, and long = average lifespan > 800 days
<sup>3</sup> Diet, sample size, and age range of mice used to generate methylome data. CD = control diet; HFD = high fat diet
<sup>4</sup> One male case; see Table S1 for individual level data
<sup>5</sup> One case excluded due to uncertain identity

### Strain-dependent patterns in global features of the methylome

Following genome-wide MBD-sequencing, we retained a set of 368,300 regions, each 150 bp in length, with sufficient coverage in the 70 samples. The majority of the CpG regions (83%) contained no sequence variants (SNPs or small insertions/deletions) segregating in the BXDs. For the 17% (62,422) with sequence variants, there was an average of 2 ± 1.6 variants within the 150 bp bin. Consistent with the DNA enrichment and filtering protocols, the 368,300 CpG regions were enriched in annotated gene features such as UTR, introns, exons and CpG islands, and also rRNA and LTRs compared to the background genome (**Table S3**).

We started with a principal component analysis (PCA). A plot of the top principal components, PC1 and PC2 (captured 19% and 13% of the variance, respectively), showed clustering of samples by strain identity, irrespective of diet (**Fig. 2a**). The one exception was a BXD65 on HFD that plotted away from the BXD65 cluster, and due to its questionable identity, it was excluded from downstream analyses, and remaining statistical tests were done in 69 samples. Sub-strains (e.g., BXD73/BXD73b; BXD65/BXD65b) also clustered in close proximity. Unsupervised hierarchical clustering confirmed the clustering of samples by strain identity rather than age or diet groups (QC plots in **Fig. S1)**

**Fig. 2.**
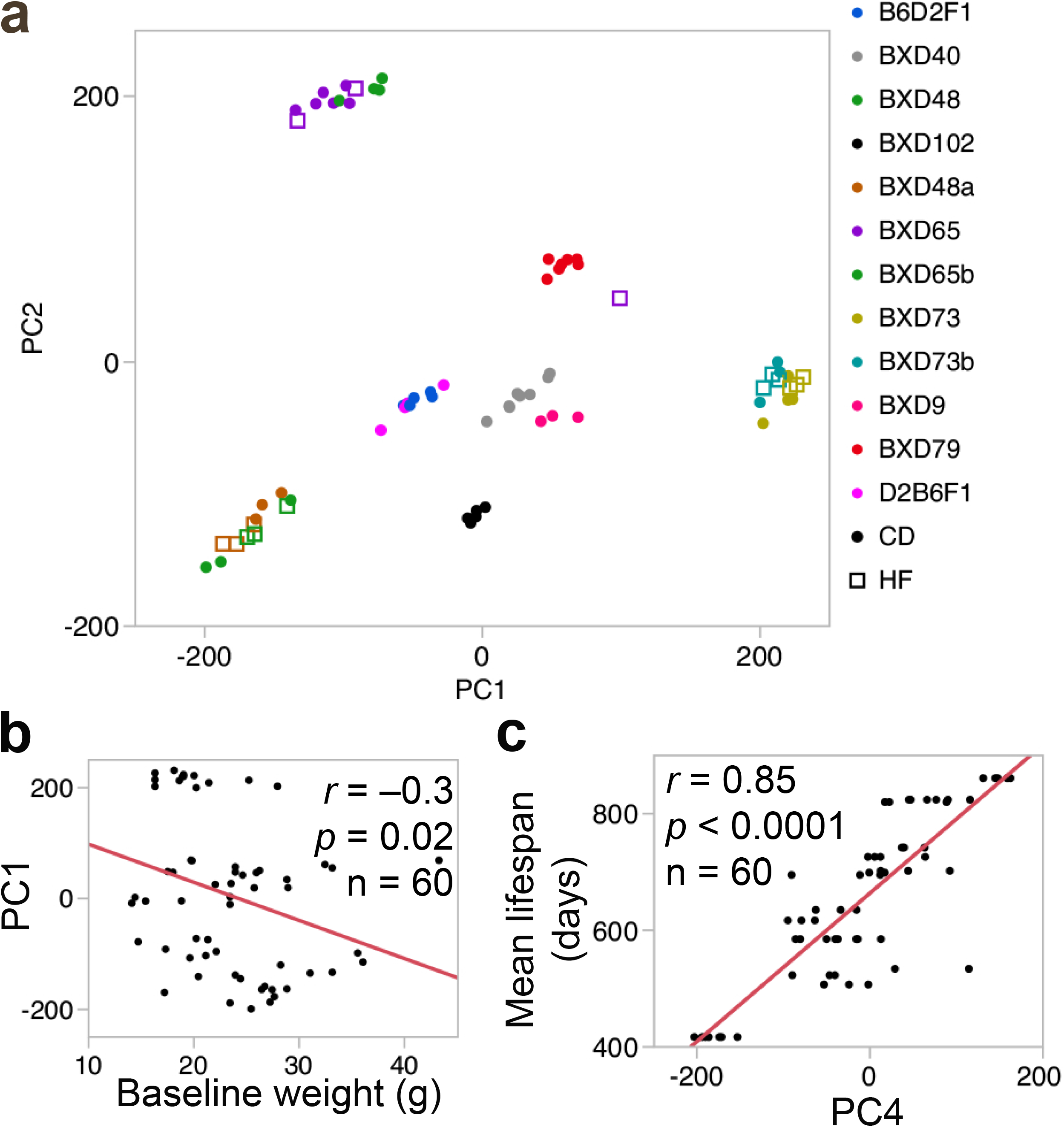
Global features of the methylome. **(a)** Scatter plot between the top 2 principal components—PC1 (19% of variance) and PC2 (13% of variance)—shows a strong population structure with mice clustering by strain identity (color coded). For strains with animals from both standard chow (CD; solid circles) and high fat diet (HFD; squares), mice on HFD co-cluster with the CD mice. **(b)** Body weight at young adulthood has a significant negative correlation with PC1. **(c)** The methylome in turn may be predictive of lifespan, and PC4 is strongly correlated with lifespan.

The top 5 PCs collectively explain 58% of the variance (**Table S1),** and we examined if these were associated with age, diet, body and organ weights, and strain lifespan. Age was not a significant correlate of any of the top 5 PCs. For strains with matched CD and HFD cases, the PCs did not differentiate between the two diets. For the weight measurements, PC1 was a significant negative correlate of BW0 (**Fig 2b**), and final body and liver weights (correlations with and without the F1s in **Table S4**). None of the other PCs were associated with the weight measurements. For the strain longevity data, PC4 (8% of variance) was the strongest correlate of mean, median, minimum, and maximum lifespan (**Fig 2c; Table S4**). PC2 and PC3 (11% of variance) were also significantly correlated with maximum lifespan (**Table S4**).

We computed the overall mean methylation for genic regions (i.e., CpG regions that overlap annotated gene features; 200,531 CpG bins in total), and intergenic regions (167,769 CpG bins), to examine whether these global features would explain the variance captured by the top PCs. Global methylation was highly strain specific (**Fig S2a, S2b; Table S1** for individual level data), and the methylation of intergenic CpGs was significantly correlated with PC1 and PC3 (**Fig S2c, S2d**). Mean methylation of genic CpGs was significantly correlated with PC4. However, mean methylation values were not directly correlated with the longevity traits. For the weight measurements, only the mean methylation of intergenic sites showed a significant positive correlation with liver weight (**Table S4**).

Taken together, the methylome-wide analysis showed that the PCs capture strain-dependent differences in overall mean methylation. These PCs in turn were associated with body weight, and strain-level lifespan. Age and diet were however not associated with the large-scale methylome features.

### Characterizing differentially methylated CpG regions

We next performed epigenome-wide tests to identify CpG regions that were associated with age, BW0, and strain median lifespan using a multiple linear mixed-model. Although aging did not have a generalized impact on global DNAm features, regression tests showed that aging was a significant predictor of site-specific DNAm at few CpG regions (**Fig 3a**). For the main effect of age, 26 regions, covering 319 CpGs, were above the 10% Bonferroni threshold (unadjusted *p* ≤ 2.7 x 10^-7^). These strong age-dependent differentially methylation regions (age-DMRs) were mostly located within genes, and 25 of the 26 bins were associated with increased methylation with age (age-hypermethylation; **Table S5**). Although BW0 and the lifespan traits had strong associations with global DNAm patterns, after partly accounting for the strain-dependent effects with the mixed model, only a few CpG regions were significant at the 10% Bonferroni threshold (**Fig 3b, 3c**). For the main effect of BW0, only seven CpG regions were significant, and all these BW0 associated differentially methylation regions (BW0-DMRs) had lower methylation among mice with higher BW0 (negative regression estimates; **Table S6**). For the strain-level median lifespan, only three CpG regions were significant at the same 10% Bonferroni threshold (**Table S7**).

**Fig. 3.**
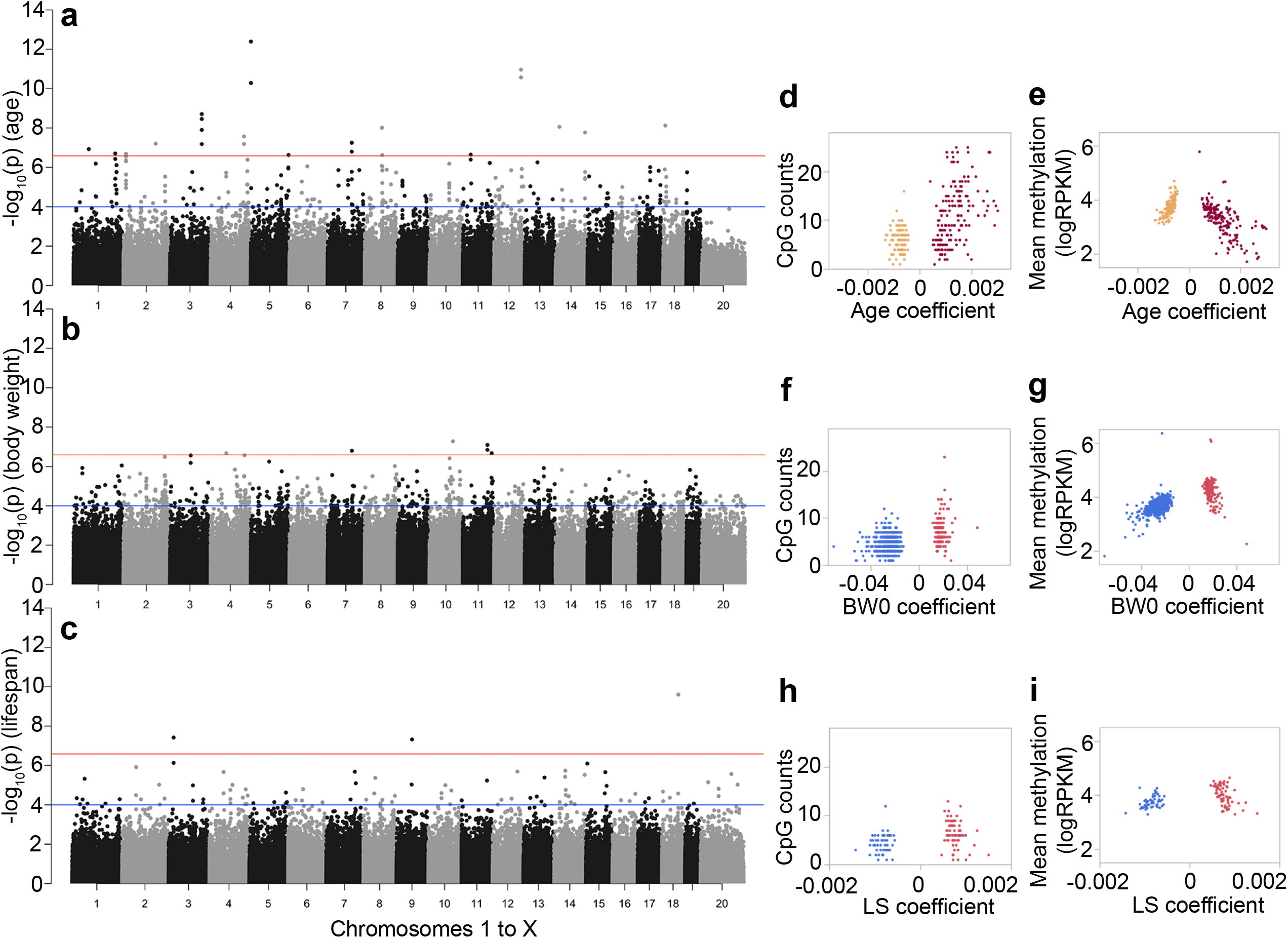
Features of differentially methylated CpGs regions (DMRs) Each point in the Manhattan plot represents the location of a CpG region (x-axis: autosomal chromosomes 1 to 19, and chromosome X as 20), and the association -log_10_*p* (y-axis) for **(a)** age effect, **(b)** body weight at young adulthood (BW0), and (c) median lifespan (LS). The genomewide significant threshold is set at -log_10_(2.7e-7) (red line; 10% Bonferroni threshold for 368,300 tests) and the suggestive threshold at -log_10_(1.0e-4) (blue line). For the age-DMRs, the regression coefficients for age (i.e., change in DNA methylation per unit change in age in days, log_10_ scale), and whether a site gained (positive coefficients, burgundy) or lost methylation (negative coefficients; sandy brown) was highly dependent on **(d)** the CpG density, and **(e)** mean methylation. For BW0-DMRs, the body weight associated coefficients also showed correlations with the CpG counts within the bin **(f)**, and sites that were negatively associated with BW0 (blue) had lower mean methylation than the sites that were positively associated with BW0 (red) **(g)**. For the LS-DMRs, whether a site was positively (red) or negatively (blue) associated with strain median lifespan was only modestly dependent on **(h)** the CpG counts, and not dependent on **(i)** the mean methylation levels.

Given the non-independence of adjacent CpG regions, we then applied a lenient threshold of uncorrected p ≤ 1.0 x 10^-4^ to define the general characteristics of the sites associated with age, BW0, and lifespan. In total, 306 CpG regions were age-DMRs at this suggestive threshold (**Fig 3a; Table S5**). A few of the age-DMRs were weakly associated with BW0 or lifespan; for instance, the top 5 age-DMRs were all age-hypermethylated and also had weak negative associations with median lifespan (p ≤ **0.05; Table S5**). However, for the most part, the age-DMRs had only modest associations with BW0 and lifespan. Of the 306, 57% were age-hypermethylated (**Table 2**). Compared to the background set of 368,300 CpG bins, the age-DMRs were highly enriched in genic regions such as promoters and exons, and CpG island, and depleted in intergenic regions (enrichment and depletion p-values in **Table S3**). For each age-DMR, we computed the average methylation and CpG density, and compared these to the age regression coefficients, which convey the change as a function of age. The regression estimates appeared to be highly dependent on local genomic characteristics, and the most pronounced changes involved age-hypermethylation (positive regression coefficients) in bins with high CpG density (**Fig 3d)** and lower average methylation (**Fig 3e**). In contrast, age-DMRs that lost methylation with age (age-hypomethylated; negative regression coefficients) featured lower CpG density, and higher average methylation (**Fig 3d, e**). Functional annotations of these regions identified significant enrichment in gene sets involved in establishment of cellular polarity (e.g., *Wnt5a, Lrrd1, Ptk7*) (**Table S8**). We consulted the human GWAS catalogue to identify genes represented by the age-DMRs that have been significantly associated with human aging and longevity (**Table S5**), and this identified only one gene, *Cux2^26^*.

At the suggestive threshold, 689 CpG regions were classified as BW0-DMRs, and the majority of these (517 or 75%) were negatively correlated with BW0 (**Table 2**). Introns was the most enriched gene feature (**Tables S3, S6**). Compared to the age-DMRs, the BW0-DMRs were in regions with relatively lower CpG density and had only 3 regions within CpG islands (**Fig 3f**). The regions that were negatively correlated with BW0 in particular, featured lower CpG densities and also lower mean methylation levels (3g). Overrepresented gene ontologies (GO) included regulation of protein homooligomerization, actin cytoskeleton reorganization, and GTPase mediated signal transduction (**Table S8**). We also note that that several of these genes have been associated with weight in human GWAS, including *Hmga2, Fto,* and *Ntrk2* (**Table S6**)^27–29^.

For lifespan, there were 124 LS-DMRs (**Table S7),** and a slight majority of these (59%, mostly intergenic sites) were positively correlated with higher median lifespan (**Table 2**). The LS-DMRs did not show any enrichment or depletion in gene features compared to the background set, and the LS-DMRs featured lower CpG density (**Fig 3h**). There was no difference in methylation levels between the DMRs that had positive or negative regression estimates (**Fig 3i**). The LS-DMRs were not enriched in any particular biological GO category. However, there was a significant enrichment in genes that result in premature death in single knockout mice, and this included an intronic DMR in the telomerase reverse transcriptase gene *(Tert)* (**Table S8**). Also enriched were genes related to abnormal eating behavior and a decrease in body mass index (e.g., *Igfbp3, Mc4r, Lpar1*), and genes related to abnormal mineral levels *(Calcr, Sptb, Wwox)*.

In terms of the potential effect of underlying sequence differences, we examined what fraction of the DMRs were sites that overlapped DNA variants segregating in the BXDs (**Table 2**). Since the background set has 62,422 bins with variants, the 51 variant containing age-DMRs did not represent a biased enrichment of such regions (hypergeometric enrichment *p =* 0.58). The BW0-DMRs and LS-DMRs were only slightly enriched in sequence variants (enrichment *p =* 0.02 and *p = 0.09,* respectively) (**Table 2**).

**Table 2.** Tally of differentially methylated regions

|  | Positive coefficients |  |  | Negative coefficients |  |  | Variants |
| --- | --- | --- | --- | --- | --- | --- | --- |
|  | N | Genic | Intergenic | N | Genic | Intergenic |  |
| **Age-DMR** | 175 | 146 (48%) | 29 (9%) | 131 | 94 (31%) | 37 (12%) | 51 of 306 (17%) |
| **BW0-DMR** | 172 | 93 (14%) | 79 (11%) | 517 | 368 (53%) | 149 (22%) | 137 of 689 (20%) |
| **LS-DMR** | 73 | 25 (20%) | 48 (39%) | 51 | 31 (25%) | 20 (16%) | 27 of 124 (21.77%) |

### Relating differentially methylated regions to gene expression

Fifty-two of the samples with liver MBD-seq data also had matched liver RNA-seq data, and we used this group to examine whether the DMRs were associated with gene expression. We linked the age-, BW0-, and LS-DMRs to the corresponding transcript. Of the 306 age-DMRs, 279 were paired with the corresponding mRNA. For these pairs, we tested how many of the DMRs were *cis*-correlated with the gene expression, and how many of the transcripts were also correlated with age of mice (**Table S9**). At a nominal *p* of 0.05 (|r| = 0.27), 79 age-DMRs (28% of age-DMRs linked to transcripts) were correlated with the expression of cognate genes (**Fig 4a**). The age-hypermethylated DMRs mostly showed positive correlations with gene expression such that the transcripts also showed increased expression with age (e.g., *Jak3, Amn, Tradd)*. The age-hypomethylated DMRs, on the other hand, were more likely to be negatively correlated with gene expression such that the transcripts showed an increase with age *(Nfkbia, Slit3)*.

**Fig. 4.**
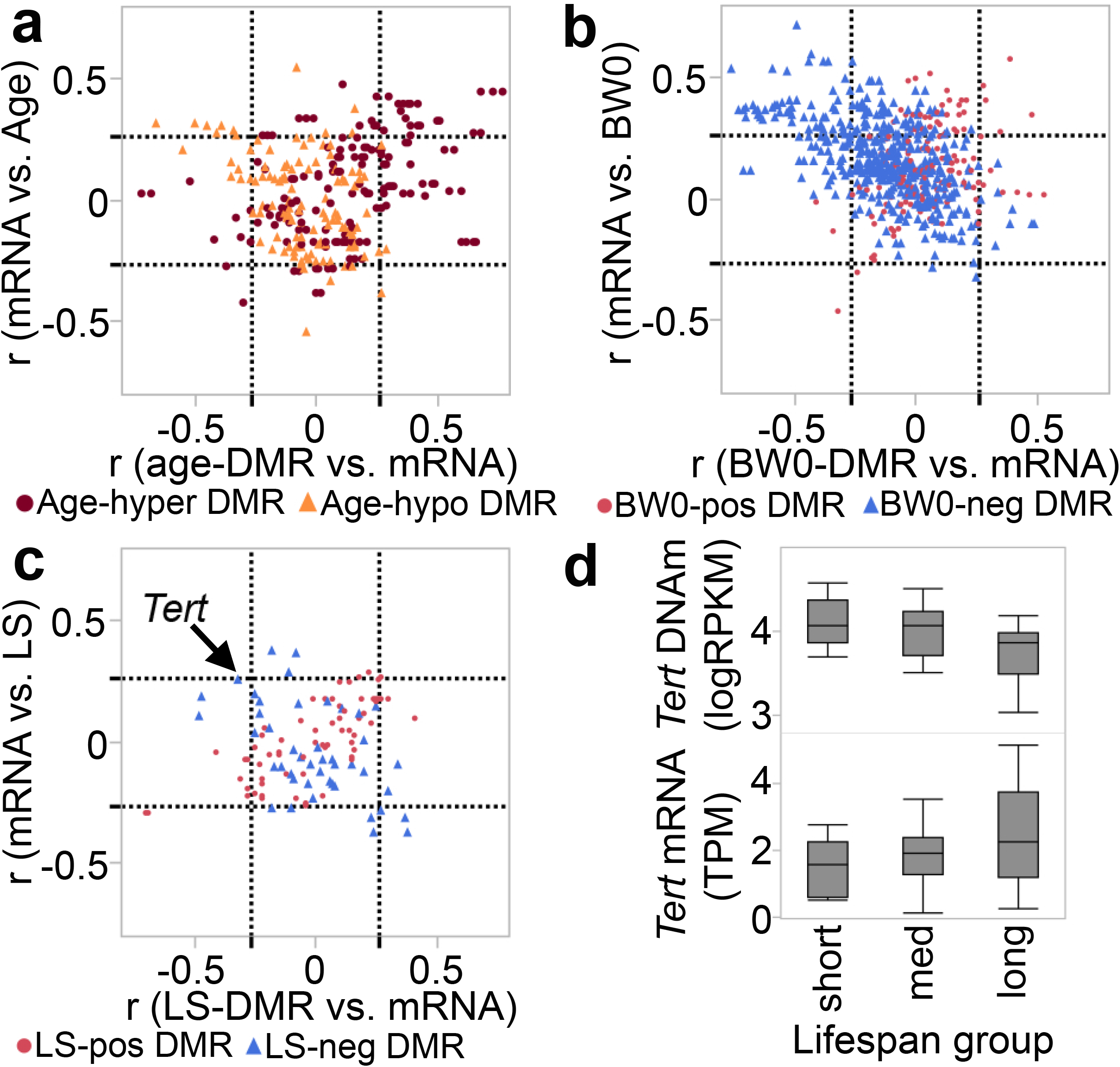
Comparison between DMRs and expression of cognate genes. **(a)** The plot shows correlations between the age-DMRs and expression of corresponding transcripts (Pearson r on x-axis), and the correlation between the transcript and age of mice (y-axis). The dashed lines demarcate the nominally significant |r| = 0.27 threshold (*p* ≤ 0.05). Most of the age-hypermethylated DMRs (burgundy circles) are positively correlated with mRNA, and these mRNAs also tend to be positively correlated with age (top right square of graph). A few of the age-hypomethylated-DMRs (sandy brown triangles) have negative correlation with gene expression, and the corresponding transcripts are positively correlated with age (top left square). **(b)** Majority of the BW0-DMRs are negatively associated with BW0 (blue triangles), and most of the DMR-mRNA pairs are located in the top-left square of the plot, i.e., gene expression is negatively correlated with DNA methylation, and positively correlated with BW0. A few of the BW0-DMRs that have positive associations with BW0 (red circles) also have positive correlations with gene expression (top right square). **(c)** The few LS-DMRs have modest correlations with gene expression. The DMR for *Tert* is negatively correlated with transcript (arrow), and the transcript has modest correlation with median lifespan at r = 0.26 (*p* = 0.07; n = 51 samples with matched MBD-seq and RNA-seq). **(d)** Comparison of DNA methylation levels (top) in the three lifespan groups shows significantly lower methylation in the long-lived group (F_2,66_ = 7.67, *p* = 0.001). A similar comparison for gene expression shows a significantly higher gene expression in the long-lived group (F_2,49_ = 3,31, *p* = 0.04).

**Fig. 5.**
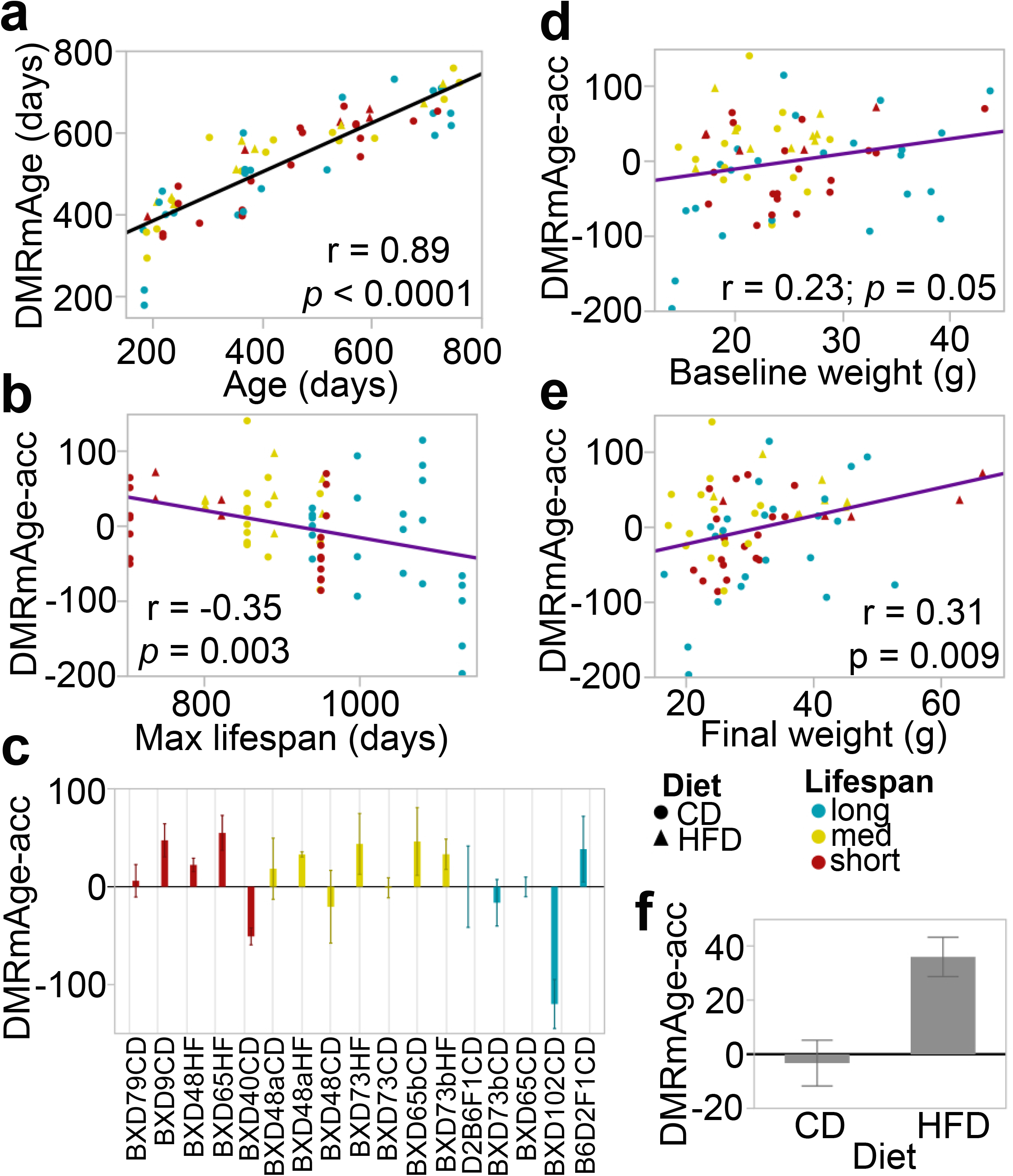
Age-DMR based measure of epigenetic aging. **(a)** The epigenetic age of mice was estimated by taking the weighted averages of the 306 age-DMRs. These age estimates, referred to as DMRmAge, has a strong positive correlation with the chronological age of mice. **(b)** The age acceleration residuals (DMRmAge-acc) derived from this clock has a significant negative correlation with the maximum lifespan data for the 17 strain-by-diet groups. **(c)** The bar plots show the mean DMRmAge-acc values for each strain-by-diet group (error bars are standard error) with the graph ordered by increasing mean lifespan (x-axis). The DMRmAge-acc is positively correlated with both body weight at young adulthood **(d)**, and final body weight **(e**). **(f)** In the BXD strains with matched samples from both control diet (CD) and high fat diet (HFD), the DMRmAge-acc is significantly higher in the HFD group compared to the CD group (−3.22 ± 36.99 in CD, 36.13 ± 27.32 in HFD, *p* = 0.002, n = 33).

For the BW0-DMRs, 614 of the CpG regions were paired to corresponding transcripts. At |r| = 0.27 (*p* ≤ 0.05), 121 (~20%) BW0-DMRs were correlated with gene expression (**Table S10**). The overall pattern indicated that the DMRs that had lower methylation in mice with higher BW0 (i.e., negative regression estimates) tended to be negatively correlated with gene expression such that there was higher expression of these genes in mice with higher BW0 (**Fig 4b**). This included *Mc5r, Nfkb1*, and *Tcf4*. Few of the DMRs that had positive associations with BW0 were negatively correlated with gene expression, and the corresponding transcripts were also upregulated in mice with higher BW0 (e.g., *Aldh18a1, Madd, Rap2a*).

For the LS-DMR, 111 paired to corresponding transcripts, and of these, only 19 CpG regions were c/s-correlated with gene expression (**Fig 4c; Table S11**). The strongest c/s-correlations were between the LS-DMRs linked to *Hpse* and *Dapp1* (r = −0.70 for both). Both CpG regions had positive regression values for lifespan, and the corresponding transcripts had lower expression in mice belonging to strains with longer lifespan. The LS-DMR in *Tert* was also negatively correlated with gene expression (**Fig 4c,** r = −0.32, *p =* 0.02), and the expression of *Tert* had a modest positive correlation with lifespan (r = 0.26, *p =* 0.07). Comparison of *Tert* methylation and gene expression among the three lifespan groups showed that the longer lived strains had significantly lower methylation levels compared to both the medium- and shortlived groups (F_2,66_ = 7.67, *p* = 0.001), and the long-lived mice had significantly higher gene expression compared to the short-lived mice (F_2,49_ = 3,31, *p =* 0.04; **Fig 4d**).

### Measure of age-acceleration from age associated CpG regions

We next evaluated whether we could derive a measure of differential rate of aging from the age-DMRs that could be predictive of lifespan differences. We first summarized the age-dependent changes by computing the weighted averages for the set of 306 age-DMRs. The weighted averages, as expected, had a strong correlation with the chronological age of mice, and for a more direct comparison, the values were scaled to the age range for the 69 samples (**Table S1**). We refer to this DMR-based age estimate as DMRmAge, and this showed a nearly linear correlation with chronological age at *r* = 0.88 (*p* < 0.0001, n =69; **Fig 4a**). Age acceleration was derived as the residuals from the regression of DMRmAge on chronological age^7,9^, with positive values indicating an older or accelerated biological age, and negative values indicatinga decelerated biological age, and we refer to this as DMRmAge-acc (**Table S1**). The DMRmAge-acc showed a significant negative correlation with strain maximum lifespan (r = −0.35, *p = 0.003,* n = 69; **Fig 4b**). It was also negatively correlated with mean and median lifespan, but this was not statistically significant (r = −0.13, *p =* 0.28 for both mean and median lifespan). The correlation between DMRmAge-acc and maximum lifespan remained significant when the analysis was limited to only the CD mice (r = −0.30, *p =* 0.03, n = 55). There were notable strain differences in average DMRmAge-acc (**Fig 4c**). These indicate that mice belonging to strain-diet groups with lower maximum lifespan have an accelerated rate of aging. We note that BXD65, which was the strain with the largest reduction in lifespan on HFD (**Fig 1a)** also showed a much higher age acceleration in the HFD group (mean DMRmAge-acc = 54.94) compared to the CD group (mean DMRmAge-acc = −0.17).

The DMRmAge-acc showed a significant positive correlation with both BW0 (r = 0.23, *p = 0.05;* **Fig 4c)** and final body weight of mice (r = 0.31, *p = 0.009)* that suggests more accelerated aging with higher body weight. Limiting to only the CD mice (n = 55), the correlation with BW0 became slightly stronger (r = 0.29, *p = 0.03),* but the correlation with final weight became weaker (r = 0.23, *p = 0.09)*. This suggests that higher body weight at baseline is associated with accelerated aging. We tested the effect of diet on the DMRmAge-acc in the BXDs that had matched samples from both diets (n = 33), and this showed a significantly higher age acceleration in the HFD group (36.13 ± 27.32 in HFD, −3.22 ± 36.99 in CD, *p =* 0.002; **Fig 4d**).

Taken together, the analysis demonstrates three points. First that the age-DMR based estimates of age acceleration is predictive of strain dependent differences in lifespan; second, that higher body weight at younger age is associated with more accelerated aging; and third, HFD significantly accelerates aging, as measured by DMRmAge-acc.

## Discussion

### Genomic features of differentially methylated regions

In the present study, we parsed the variance in the methylome and defined site-specific methylation differences that may be attributed to a strain-level phenotype, median lifespan, and two individual-level variables—age and BW0. For the age-DMRs, the time-dependent patterns were consistent with previous reports^14,30^. As in Sziráki et al.^14^, methylation loss over time occurred mostly in regions with higher average DNAm, and methylation gains occurred mostly in regions with lower average DNAm. Since DNAm was quantified over 150 bp nonoverlapping bins, we were also able to relate the differential methylation patterns to the local CpG density. In particular, for the age-hypermethylated DMRs, the increase in DNAm with aging had a strong positive correlation with CpG density. This too, is consistent with reports that CpG dense regions—a feature of CpG islands, which typically remain unmethylated—are the sites that tend to gain methylation with age^31^. The genes represented by the age-DMRs included a few notable members such as *Cyp46a1* and *Abca7,* which are involved in cholesterol metabolism and implicated in Alzheimer’s disease^32^, and few members of the WNT-signaling and mesenchyme developmental pathways (e.g., *Fzd1, Fzd8, Wnt5a, Jak3, Ptk7, Nrp2*). The current data replicated the CpG islands in *C1ql3* and *Ptk7*, which we previously reported as age-hypermethylated sites in the BXD parental strains, B6 and D2^33^. A region-based functional annotation analysis revealed a significant enrichment in genes involved in cell polarity, an aspect of cells that is established during development through interaction with the WNT polarity signaling pathway, and which becomes dysregulated during aging^34,35^. In terms of the potential impact on gene expression, the general pattern was for the cognate genes to increase in expression with age regardless of whether the corresponding CpG regions were age-hypomethylated or age-hypermethylated.

Body weight, even at young adulthood and before introduction to HFD, was significantly associated with the global methylome patterns, and was also predictive of strain longevity. We therefore included BW0 as a predictor variable in the multiple regression model. BW0 was associated with 689 CpG regions, most at the suggestive level, and these BW0-DMRs were significantly enriched in GO categories related to protein assembly, rho and GTPase mediated signaling, and oligosaccharide metabolism. While genes related to fat metabolism or adiposity were not enriched, we note that there were a few genes that have been previously associated with body weight by human GWAS at p < 5e-08^26^. This included an intron DMR in the fat mass and obesity associated *Fto* gene, which plays a key role in energy homeostasis, and consistently shown to influence body weight in humans that may be mediated by differences in DNAm and gene expression^36^. The BW0-DMRs were mostly located within genes, particularly introns, and although the differential methylation patterns showed some relation with the local CpG counts, this was not as pronounced as in the case of the age-DMRs. The more striking aspect of the BW0-DMRs was that 75% were negatively associated with BW0. For the BW0-DMRs that were linked to the corresponding transcript, the overall pattern indicated a negative correlation between DNAm and gene expression. This means that while DNAm at these sites were, in general, inversely correlated with body weight, the transcript levels of the corresponding genes generally had higher expression in mice that were heavier.

For the longevity trait, only 124 CpG regions were significant at the suggestive threshold. These sites were not enriched in any particular genomic feature or GO category, and did not show a strong dependence on CpG density or mean methylation. A point of distinction for the LS-DMRs is that, while the age- and BW0-DMRs were related to individual-level variables, the LS-DMRs were related to life expectancy based on the strain and the diet. This indirect association with the phenotype may explain why only few DMRs were uncovered by the current mixed model. Given this small number of genes, it is particularly striking that the region-based annotation revealed that genes that cause premature death in single knockout mice are the most enriched gene set among the LS-DMRs (phenotype ID MP:0002083, defined as “death after weaning, but before the normal lifespan” in http://www.informatics.jax.org). For these functionally critical genes that result in premature death when deleted, our results imply that quantitative differences in DNAm at these sites can modulate strain variation in normal lifespan. Among the LS-DMRs, both *Tert* and *Igfbp2* have also been linked to aging and longevity by human GWAS^37,38^.

### Building clocks from age dependent CpG regions

Currently, there are several different versions of the DNAmAge estimator available for both mice and humans, some optimized for a specific tissue, and others with multi-tissue application^2,3,5–8^. The standard protocol for developing DNAmAge clocks starts by applying a regression algorithm in a training dataset, followed by age estimation in validation cohorts to gauge the accuracy of the clock. Our goal in this study was not to develop another DNAmAge clock. Given the sample size of the present study, that would not have been a feasible pursuit. Instead, our goal was to test whether the age-dependent CpG regions could discern lifespan differences between the mouse strains, and how the rate of epigenetic aging relates to body weight and diet. For this, we simply summarized the age-DMRs by computing the weighted averages for each sample. The weighted averages, unsurprisingly, correlated strongly with the chronological age of mice. More importantly, the age acceleration derived from the age-DMR based clock was (a) inversely correlated with the strain lifespan phenotype, (b) was significantly more accelerated in the HFD group, and (c) was positively correlated with body weight. This conveys that the age-DMRs can estimate genetically modulated differences in rates of biological aging and lifespan, and is a modifiable outcome that is accelerated by HFD. Furthermore, the results highlight the interdependence between body weight, diet, and health and aging, and our observations agree with the well-known influence of body mass on longevity, and the more favorable health profile associated with lower body mass^39^.

The strain with the most decelerated clock, and presumably slowest rate of biological aging, was BXD102 on CD, which is also the longest-lived BXD strain we had in the study (**Table 1**). For these mice, the DMRmAge-acc for the females ranged from −78 to −196 days. Interestingly, the one male BXD102 had the least decelerated clock among the BXD102 samples at −66 days. The most accelerated epigenetic aging was seen for the short-lived BXD9 on CD, and BXD65 on HFD. However, we note that there were a few mismatches between strain lifespan classification and the DMRmAge-acc. BXD40 on CD, although classified as short-lived, was the only short-lived strain with a mean negative DMRmAge-acc. The F1 hybrids, on the other hand, although classified as long-lived, were the only long-lived groups with mean positive DMRmAge-acc. Unlike the inbred BXDs, the heterozygous F1s have hybrid vigor in both body weight and lifespan, and this likely explains the inconsistency.

### Technical considerations and caveats

Before concluding, we should address a few caveats. The sequence alignment was done to the mm10 B6 reference genome, which means that for regions with genetic variants segregating in the BXDs, the sequence differences could compromise alignment. The strong “population stratification” in the PC plot, reminiscent of human genetic data, is likely the result of true quantitative variation in methylation, and also due to a portion of the CpG regions serving as surrogates for underlying genotype^40^. Aging is independent of genetic background and the age-DMRs are expected to be less susceptible to the confounding effect of DNA sequence variants. For body weight and lifespan, the interpretation is complicated by the fact that both phenotypes are closely linked to genotype, and the DMRs may reflect true differences in DNAm levels, or differential quantification due to sequence effects. To partly control for this, we used a mixed model that fitted each strain-diet group as a random intercept. For the age-DMRs, 17%of the CpG regions contained sequence variants, and this is similar to the 17% of variant containing bins in the background set of 368,300 regions. The BW0-DMRs and LS-DMRs were only slightly enriched in variant containing bins, and for the most part, the CpG regions were devoid of sequence differences.

To our knowledge, almost all existing DNAm clocks have relied on bisulfite-based assays, which have the advantage of providing single CpG resolution^2–8^. For the mouse model, the reduced representation data (RRBS) generated by Petkovich et al.^5^ from ~141 B6 mice, has been used in different studies to define tens of thousands of age dependent CpG sites (over 43,000 CpGs in Lowe et al.^41^, and over 146,000 in Szirki et al.^14^). Compared to these, the present work identified only 306 age-DMRs at the suggestive threshold. These CpG regions cover 2,691 CpG sites, and this is still a relatively modest number of age-dependent CpGs. This is likely a result of the smaller sample size, and the rigorous statistical model due to the different genotypes that constituted the present cohort. Another contributing factor may be the methylome assay we used, as MBD-seq provides lower quantitative sensitivity than the bisulfite based assays^42^. Nonetheless, MBD-sequencing still delivers highly sensitive and replicable quantification of genome-wide methylation^25^. Using alternate assay methods also demonstrates that the DNAm age estimators are robust to the techniques used to quantify DNAm.

## Conclusion

In conclusion, our results demonstrate that the epigenetic clock defined from age-DMRs is sensitive to subtle differences in natural lifespan that arise from common genetic variants, and is modifiable by environmental interventions, such as diet. The intercorrelations between epigenetic aging, body weight, and longevity also provide evidence that the methylome could provide a mechanistic link between the well-known effect of body mass on aging and lifespan.

## Experimental Procedures

### Statistical analysis

Descriptions of sample processing for MBD-seq, sequence alignment, initial bioinformatics, and data filtering are provided in **Appendix S1**. In brief, DNAm quantification was done by counting sequenced reads at every 150 bp non-overlapping bins. After removing bins with low coverage, we retained 368,300 CpG regions for further analysis. PCA and hierarchical clustering was done in R for this full set of bins (**Fig S1a**). These detected no outlier samples and the DNAm profiles were consistent across the samples (**Fig S1b)**, and averaged at 3.8 ± 0.72 logRPKM. The CpG regions were then divided into bins that occurred within annotated genes (genic set, 200,531 bins), and those that were in intergenic regions (167,769 bins). For each sample, the overall average methylation and variance for these genic and intergenic sets were computed. The intercorrelations between the large-scale methylome features, body weight measures, and strain-level lifespan phenotype were examined using Pearson correlations.

To detect DMRs, we applied the following model using the lme4 R package (v4_1.1-21)^43^: Imer(logRPKM ~ age + BW0 + medianLifeSpan + (1| StrainDiet)). Manhattan plots were generated using the qqman R package^44^. For the DMRs, we performed two types of enrichment analyses. First, we evaluated relative enrichment in genomic features (i.e., introns, exons, CpG islands, etc.) compared to the background set using the hypergeometric test in R (R codes in **Table S3**). Following that, we carried out functional annotation and enrichment analysis against the whole background genome using the GREAT application (version 4.0.4)^45^. We followed the GREAT recommendation of calculating enrichment using both a binomial test and a hypergeometric test, and reporting only categories that are significant by both methods (FDR < 0.05). To search for human GWAS hits, we referred to the GWAS catalogue^26^, and searched for the terms “body weight”, “aging”, and “longevity”. For variants with reported association *p ≤* 5.0e-08, the mapped human gene symbols were then matched to the corresponding mouse genes associated with the DMR.

### Estimating epigenetic age and age acceleration

To estimate epigenetic age, we computed the weighted averages with each of the 306 age-DMRs weighed by the respective age regression coefficient. The weighted averages were then scaled to the age range in the 69 samples using the following formula: DMRmAge = {(((weighted-average – min.weighted.average) x age.range) / weighted.average.range) + min.age}, where min.weighted.average and weighted.average.range are the minimum value and range for the weighted averages in the 69 BXD samples, and age.range = 578 days is the range of chronological age in the 69 BXDs, and min.age = 181 days is the minimum age for the 69 BXDs. As recommended in Thompson et al. 2018^7^, the “age acceleration” was computed as the residuals after fitting the predicted age to chronological ages: residuals(lm(DMRmAge ~ Age)).

### Transcriptomes analyses

We used liver gene expression data that are available from GeneNetwork (http://genenetwork.org; data accession ID GN877, UTHSC BXD Harvested Liver RNA-Seq (Oct19)). While this is a bigger cohort, we limited the analysis to only 52 mice that had matched MBD-seq in the present study. Gene expression was log_2_ transformed transcripts per million (TPM). For the age-, BW0-, and LS-DMRs, the cognate genes were defined as the gene in which the DMR is located for genic sites, or the nearest gene promoter for intergenic regions. These were then matched by gene symbol to the corresponding transcript. For *cis*-correlations, we performed a Pearson correlation between the DNAm levels and transcripts levels. In cases where a DMR paired to multiple transcript variants, we retained only the DMR-mRNA pair that had the strongest *cis*-correlation such that each DMR was paired to only a unique mRNA. Following this, we used Pearson correlations to relate the expression levels with the corresponding variable, i.e., age, BW0, or median lifespan.

## Acknowledgements

The present work relied on resources from the BXD Aging Colony, and we thank the entire team, particularly Dr. Lu Lu, Jesse Ingels, Casey J Chapman, Melinda S McCarty, Arthur Centeno, and Dr. Megan K Mulligan. We thank Dr. Saunak Sen for his invaluable comments on the statistics. We thank the UTHSC-Rhodes College Population Health Researcher Program and Dr. Teresa Waters for support and for bringing an excellent summer student to the PI’s lab. We thank Dr. Karolina A. Aberg and her team at the Virginia Commonwealth University for their advice on the MBD-seq technique. This study was funded by the NIH NIA grants R21AG055841 and R01AG043930.

## Author contribution

KM supervised the study, contributed to conception of study design and analysis, and wrote the manuscript. RWW is the principal investigator of the BXD Aging Colony and contributed to the conception of the aging study, and provided access to the biorepository resource. JVSS, AHBH, and EGW contributed to the lab work, and JVSS performed the primary bioinformatics and initial data processing. DA and SR contributed to analysis and data acquisition. All authors contributed to and approved the final version of the manuscript.

## Data availability

The normalized MBD-seq data for the 368,300 CpG bins (including chromosomal coordinates, region annotations, CpG density, and variant counts) are available from the NCBI NIH Gene Expression Omnibus. The full raw fastq files are available from the NCBI NIH Sequence Repository Archive. GEO accession ID is GSE137277, and the linked SRA accession ID is SRP221380. (currently set to private but will be made available upon official publication)

Ethics approval. All animal procedures were in accordance to protocol approved by the Institutional Animal Care and Use Committee (IACUC) at the University of Tennessee Health Science Center.

