## Supplemental figures for "Body weight at young adulthood and association with epigenetic aging and lifespan in the BXD murine family"

**Supplementary Figures**

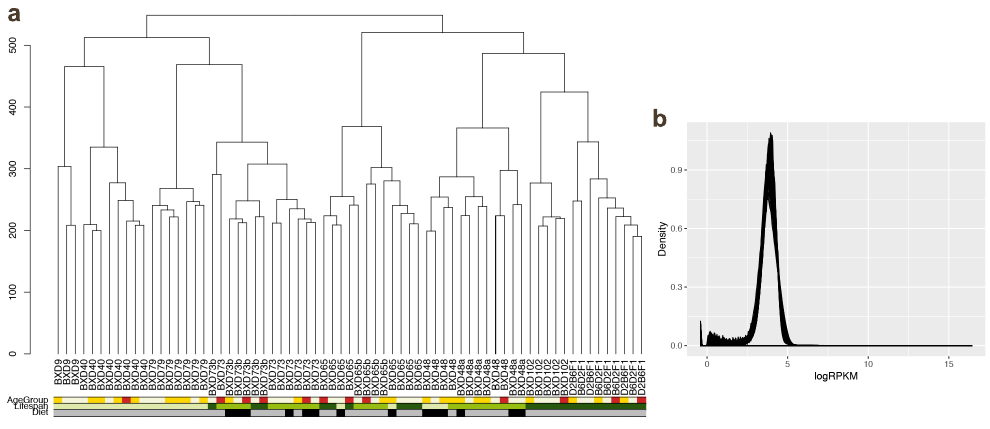

**Fig S1. Unsupervised hierarchical cluster and density plots**

**(a)** The dendrogram plotted using the genome-wide data shows that samples cluster by strain identity rather than age or diet. **(b)** Density plots of logRPKM values for the 368,300 CpGs bins show a consistent distribution. Methylation scores average is at 3.8 ± 0.72 logRPKM.

**Fig S2. Genome-wide mean methylation and correlations with principal components**

For each individual mouse, the overall mean methylation and within-individual variance was calculated for **(a)** 167,769 CpG regions in intergenic sites, and **(b)** 200,531 genic CpG regions located within annotated genes. The intergenic regions have wide variation between strains and the F1 hybrids have the highest mean methylation and lowest variance. The genic CpG regions are more consistent across strains. Mean methylation is inversely correlated with variance, and this is particularly pronounced for the intergenic CpG regions. Average methylation at intergenic regions (x-axis) is correlated with PC1 **(c)**, and PC3 **(d)**, and average methylation at genic regions is correlated with PC4 **(e)**.
