## Appendix for "Body weight at young adulthood and association with epigenetic aging and lifespan in the BXD murine family"

**Appendix S1**

**Sample preparation**

Liver samples were obtained from the BXD colony maintained at the University of Tennessee Health Science Center (UTHSC; see Roy et al. ^1^). All animal procedures were in accordance to the protocol approved by the UTHSC Institutional Animal Care and Use Committee (IACUC). DNA was extracted using the DNeasy Blood & Tissue Kit from Qiagen. Nucleic acid purity was inspected by using a NanoDrop spectrophotometer, and quantified using a Qubit fluorometer dsDNA BR Assay. Affinity-based enrichment was carried out using the MethylMiner DNA enrichment kit from ThermoFisher Scientific according to the manufacturer’s standard protocol. In brief, 1 μg of DNA in 110 μl of low TE buffer was fragmented to ~150 bp using a Covaris S2 ultrasonicator. Sonication settings were: cycle/burst of 1 for 10 cycles of 60 s, duty cycle of 10%, intensity of 5.0. DNA fragment size and quality were assessed using the Agilent Bioanalyzer 2100. Following MBD capture reactions, DNA was eluted in a single step using high salt (2000 mM NaCl) elution buffer, and re-concentrated by ethanol precipitation. The final concentration of methylated-CpG enriched DNA ranged from 1.12 to 5.43 ηg per μl (2.46 ± 0.94). Sequencing was done to a depth of approximately 50 million reads per sample (150 bp paired-end) on the Illumina HiSeq 4000, and carried out by Novogene (<https://en.novogene.com>).

**Read alignment and initial data processing**

FASTQ files were first inspected with the FastQC tool (v.0.11.8) ^2^, and alignment was done to the mouse reference genome (mm10/GRCm38) using Bowtie2 (v.2.3.4.3) ^3^. Alignment quality was evaluated with SAMtools (v.1.9) ^4^, and SAMstat v.1.5.1^5^. Potential PCR duplicates and reads with mapping quality less than 10 were removed. The bam files were loaded to the MEDIPS R package (v.1.36.0) ^6^ for additional quality checks and assessment of read coverage. Saturation analysis (MEDIPS.saturation) showed that all 70 libraries had sufficient read coverage, and pair-wise correlations (MEDIPS.correlation) showed high consistency between samples (Pearson *r* > 0.90 for all pairs). Given the MBD enrichment, all the samples were enriched for CpGs (mean CG enrichment score of 2.30 ± 0.19; using the MEDIPS MEDIPS.CpGenrich function), and on average, 51% of CpGs in the reference genome was covered by at least one mapped read, with 28% of CpGs covered by > 5 mapped reads. To quantify DNAm, the mouse genome was divided in 150 bp non-overlapping windows and reads were counted for each bin with normalization to the local CpG density (coupling factor or CF) using the function MEDIPS.meth and the following parameters: ws =150, extend = 150, uniq = 1, shift=0. Read counts were then filtered to retain only 150 bp bins that had sufficient coverage for reliable quantification and statistical analyses. First, bins with no CpGs (CF = 0) and mean read counts ≤ 1 were excluded, resulting in 4,286,826 bins. The Y chromosome was also excluded. The filtered data was loaded to the EdgeR R package (v3.24) ^7^ and further filtered on the basis of counts per million (CPM) to retain only reads with more than 1 CPM in 2 or more libraries. This resulted in 368,300 CpG regions with sufficient coverage across the libraries, and these were normalized using the calcNormFactors function. RPKM values extracted using the parameters gene.length = 150, log = TRUE. The compendium of SNPs and small insertions/deletions segregating in the BXDs have been catalogued for the BXDs ^8^, and we used this information to count the number of variants in each of the 368,300 150 bp bins. These CpG regions were annotated for genomic features using the HOMER program (v4.10) ^9^.
